# Differential TLR-ERK1/2 activity promotes viral ssRNA and dsRNA mimic-induced dysregulated immunity in macrophages

**DOI:** 10.1101/2024.05.24.595760

**Authors:** Rakshya Shrestha, Paige Johnson, Roshan Ghimire, Cody Whitley, Rudragouda Channappanavar

**Affiliations:** Department of Veterinary Pathobiology, College of Veterinary Medicine, Oklahoma State University. Stillwater, OK, 74078; Oklahoma Center for Respiratory and Infectious Diseases, Oklahoma State University, Stillwater, OK, 74078

**Keywords:** SARS-COV-2, TLRs, ERK1/2, interferon, inflammation, and macrophages

## Abstract

RNA virus induced excessive inflammation and impaired antiviral interferon (IFN-I) responses are associated with severe disease. This innate immune response, also referred to as ‘dysregulated immunity,’ is caused by viral single-stranded RNA (ssRNA) and double-stranded-RNA (dsRNA) mediated exuberant inflammation and viral protein-induced IFN antagonism. However, key host factors and the underlying mechanism driving viral RNA-mediated dysregulated immunity are poorly defined. Here, using viral ssRNA and dsRNA mimics, which activate toll-like receptor 7 (TLR7) and TLR3, respectively, we evaluated the role of viral RNAs in causing dysregulated immunity. We show that murine bone marrow-derived macrophages (BMDMs) stimulated with TLR3 and TLR7 agonists induce differential inflammatory and antiviral cytokine response. TLR7 activation triggered a robust inflammatory cytokine/chemokine induction compared to TLR3 activation, whereas TLR3 stimulation induced significantly increased IFN/IFN stimulated gene (ISG) response relative to TLR7 activation. To define the mechanistic basis for dysregulated immunity, we examined cell-surface and endosomal TLR levels and downstream mitogen-activated protein kinase (MAPK) and nuclear factor kappa B (NF-kB) activation. We identified a significantly higher cell-surface and endosomal TLR7 expression compared to TLR3, which further correlated with early and robust MAPK (pERK1/2 and p-P38) and NF-kB activation in TLR7-stimulated macrophages. Furthermore, blocking EKR1/2, p38, and NF-kB activity reduced TLR3/7-induced inflammatory cytokine/chemokine levels, whereas only ERK1/2 inhibition enhanced viral RNA-mimic-induced IFN/ISG responses. Collectively, our results illustrate that high cell surface and endosomal TLR7 expression and robust ERK1/2 activation drive viral ssRNA mimic-induced excessive inflammatory and reduced IFN/ISG responses, and blocking ERK1/2 activity would mitigate viral-RNA/TLR-induced dysregulated immunity.

## INTRODUCTION

Emerging and re-emerging RNA viruses pose a significant threat to global public health. As opposed to the only dsRNA virus (rotavirus) infecting humans, several dozens of positive and negative sense ssRNA viruses cause disease in humans. Most of these ssRNA viruses have zoonotic sources, and their accidental spillover from animal reservoirs into human population has caused multiple epidemics and pandemics ^1 2^. In addition to the viruses currently circulating in humans or periodically spilled over from animal sources, many ssRNA viruses with zoonotic potential to infect and cause outbreaks in humans have been identified in bats and several other animal reservoirs ^3 4 5 6^. Interestingly, compared to DNA viruses (e.g., herpesviruses, adenoviruses, hepadnavirus, and papillomaviruses), the majority of emerging and re-emerging ssRNA viruses cause acute and often severe disease in humans ^7^. The fatal illness caused by these RNA viruses is commonly associated with excessive inflammation and impaired antiviral immunity (dysregulated immunity) ^8 9 10^. Consequently, to better understand severe disease caused by the ssRNA viruses, it is critical to define the virus and host factors that drive excessive inflammation and impair antiviral immunity.

Viruses induce antiviral and inflammatory responses following recognition of viral pathogen-associated molecular patterns (PAMPs) by cognate immune sensors ^11 12 13^. The viral ssRNA genome and replication intermediate dsRNA are two key PAMPs that elicit antiviral and inflammatory response during ssRNA virus infections ^14 15^. Among the two viral RNA molecules, ssRNA is highly abundant, and by some estimates, the ratio of ssRNA to dsRNA is >100 to 1 in an active ssRNA virus replicating cell ^16^. Following infection, the intracellular ssRNA is sensed by retinoic acid-inducible gene I (RIG-I) receptor while dsRNA is detected by both RIG-I and melanoma differentiation-associated protein 5 (MDA-5) ^17 18 19 20^. In contrast, extracellular viral ssRNA and dsRNA, derived from a lytic infection, are detected by cell surface and endosomal located TLR7/8 and TLR3, respectively ^21 22 23^. TLR7/8 utilizes myeloid differentiation primary response 88 (Myd88) and TLR3 signals through TIR-domain-containing adaptor-inducing beta interferon (TRIF) to elicit antiviral and inflammatory responses ^21 24 25 26^. TLR-Myd88 or TRIF signaling further activates mitogen-activated protein kinase (MAPK) or nuclear factor kappa B (NF-kB) to induce inflammatory mediators and interferon regulatory factors (IRFs) to facilitate IFN/ISG response ^27 28 29 30^. However, the relative contribution of TLR7 and TLR3 activation and the mechanistic basis for TLR7- and TLR3-induced differential antiviral and inflammatory response are not clearly defined.

Viruses possess multiple structural and non-structural proteins (NSPs) that directly interact with IFN signaling proteins, ISGs, and NF-kB to subvert antiviral responses ^31 32 33^. However, the virus-induced antagonism of IFN/ISG activity without directly interacting with intracellular signaling protein(s) is poorly described. Therefore, the primary objective of this study is to elucidate the role of differential cell surface and endosomal TLR3/TLR7 expression and downstream ERK1/2 activation in viral RNA-induced dysregulated immunity. In this study, to mimic extracellular viral RNA sensing by cell surface or endosomal TLR3 and TLR7, we treated murine bone marrow-derived macrophages (BMDMs) with R848 (ssRNA mimic/TLR7 agonist) and PolyIC (dsRNA mimic/TLR3 agonist). Our results illustrate that viral ssRNA and dsRNA mimics induce differential antiviral and inflammatory cytokine responses, which correlates with cell surface/endosomal TLR3/TLR7 expression and MAPK and NF-KB activation. We also show that viral RNA-mimic-induced robust ERK1/2 activation drives dysregulated immunity. Collectively, our results highlight an under-appreciated role of viral ssRNA- and dsRNA-mediated differential ERK1/2 activation in virus-induced dysregulated immunity and, by extension, severe disease.

## MATERIALS AND METHODS

### Murine macrophage extraction and stimulation of macrophages

Bone marrow cells extracted from C57BL/6 mice were grown in 12-well plates using RPMI medium supplemented with 10% fetal bovine serum, 1% penicillin-streptomycin, 1% L-glutamine, 1% sodium bicarbonate, 1% sodium pyruvate, 1% non-essential amino acids and were treated with macrophage colony-stimulating factor (m-MCSF, GenScript, Cat # Z02930-50, 40 ng/ml) for 6-7 days with media change every three days to differentiate bone marrow cells into mouse bone marrow-derived macrophages (BMDMs).

### TLR stimulation and MAPK/NF-kB inhibitor studies

For stimulation studies, a confluent monolayer of primary BMDMs was stimulated with R848 (InvivoGen, Catalog # tlrl-r848-5, single-stranded RNA mimic) and Poly I:C (InvivoGen, Catalog # tlrl-pic, double-stranded RNA mimic) each treated at 5 or 10ug/ml for 8 or 24 hours. After 8 and 24 hours of stimulation, cells in Trizol or cell supernatants were collected to assess mRNA and protein levels, respectively, of inflammatory cytokines and chemokines and type-I interferons (IFN-Is). Primary mouse BMDMs were treated with pathway-specific inhibitors for inflammatory pathway inhibition studies, each at 5μM concentration. Specifically, we used inhibitors of ERK1/2 (Trametinib, SelleckChem, Catalog # S2673), p38 (Losmapimod, Selleckchem Catalog #S7215), NF-kB (BAY11-7082, Catalog # S2913) and JNK inhibitor (Millipore-Sigma Catalog # CAS 129-56-6) 2 hr before stimulation with R848 or Poly I:C until sample collection at 8 or 24hrs for mRNA or protein levels of antiviral and inflammatory cytokines and chemokines.

### Enzyme-linked immunosorbent assay (ELISA)

We used sandwich ELISA to quantify cytokine levels. Concentrations of TNF-α (BD Biosciences., San Diego, CA, USA Cat#558534), IL-6 (BD Biosciences., San Diego, CA, USA Cat # 555240), MCP-1 (BD Biosciences., San Diego, CA, USA Cat # 555260), and IFN-β (R&D systems., Minneapolis, MN, USA Cat #DY8234-05) in cell culture supernatants were measured using commercially available BD OptEIA™ or Bio-techne ELISA kits as per manufacturer’s instructions. A 96-well high-binding microplate (Corning, Catalog # 3361) was coated with capture antibody overnight at 4°C or room temperature. The next day, plates were blocked for 1 hour at room temperature and 100 μL of samples were added to each ELISA well and incubated for 2 hrs. Washing was performed 2-3 times at each subsequent step. Detection antibody was added to each well and incubated for 1 hour, followed by the addition of an HRP-conjugated enzyme reagent. The plates were incubated for 30 minutes in the dark using TMB substrate (BD Biosciences., San Diego, CA, USA Cat #555214). The colorimetric reaction was stopped by adding 2N H_2_SO_4,_ and absorbance was measured at 450nm wavelength using a spectrophotometer (SpectraMax® M2e).

### Quantitative reverse transcription polymerase chain reaction (qRT-qPCR)

Total RNA was extracted from BMDMs using a Trizol reagent (TRIZOL, Invitrogen Cat #15596026) as per manufacturer’s instructions. Trizol-extracted RNA was treated with RNase-free DNase (Promega Catalog: M610A) to remove any genomic DNA contamination and RNA was quantitated using Nanodrop (Nanodrop™ Thermoscientific). cDNA was synthesized using m-MLV Reverse Transcriptase (Invitrogen Cat # 28025013). Quantitative gene analysis was performed using PCR master mix (2X qPCR Universal Green MasterMix Cat # qMX-Green-25 ml, Lamda Biotech) and real-time PCR system (Applied Biosystems 7500 QuantStudio 6 Pro). The Ct value of target genes obtained from each sample was normalized to housekeeping gene GAPDH and relative gene expression was quantified using 2^-ΔΔct^ values.

### Western blot assay

Total cell extracts were lysed in RIPA buffer (Cell signaling, catalog # 9806) supplemented with protease inhibitor (Thermo Scientific., Cat # 1861281) post-RNA-mimic treatment (10, 30, and 60 min). Total protein concentrations were determined using a Pierce™ BCA protein assay kit (Thermofisher Scientific, Cat # 23225). An equal amount of protein was loaded for SDS-PAGE on 10% polyacrylamide gels, electro-transferred to nitrocellulose membrane (Bio-Rad 0.45 μM #1620115), and probed with primary antibody overnight at 4°C. Primary antibodies used are: anti-phosphorylated p44/42 MAPK (Cell signaling # 9101, 1:1000), anti-p44/42 MAPK (Cell signaling # 4695, 1:1000), anti-phosphorylated p38 MAPK (Cell signaling # 4511, 1:1000), anti-p38 MAPK (Cell signaling # 8690, 1:1000), anti-phosphorylated SAPK/JNK (Cell signaling # 9255, 1:1000), anti-SAPK/JNK (Cell signaling # 9252, 1:1000), anti-phosphorylated NF-kB p65 (Cell signaling # 3033, 1:1000), anti-NF-kB p65 (Cell signaling # 8242, 1:1000), and β-actin (Invitrogen # MA5-15739). The membrane was washed 3X times with TBST wash buffer and incubated with appropriate secondary antibody for 1 hour, then visualized using Amersham Imager 600. Secondary antibodies used are Anti-Mouse IgG (Jackson ImmunoResearch Cat #115-035-003) and Anti-Rabbit IgG (Invitrogen # 31460). Beta-actin was used as a loading control. For densitometry, images are analyzed using Image J software.

### Flow Cytometry

Surface and intracellular expression of TLR3 and TLR7 on BMDMs were determined by FACS analysis. A total of 1.5 × 10^5^ cells/well were seeded on a 96-well round bottom plate. For cell surface staining, cells were stained with PECy7 α-CD45 (clone: 30-F11, Biolegend: cat # 103114), V450 α-CD11b (clone: M1/70, Biolegend: Cat # 101224), APC α -F4/80 (clone: BM8, BioLegend: Cat #123116), PE α-TLR3 (clone:11F8, Biolegend: Cat 141904), and PE α-TLR7 (clone: A 94B10, BD Biosciences: Cat # 565557) and incubated for 20 min at 4°C in the dark. Cells were washed once with FACS buffer (1X PBS; 2% FBS, 0.05% NaA2), followed by centrifugation at 1200g for 5 min. Intracellular expression of TLR3 or TLR7 was determined following fixation and permeabilization using a Cytofix/Cytoperm Kit (BD Bioscience Catalog # 554655). All antibodies were diluted 1:200 in FACS buffer for surface staining or perm buffer for intracellular staining. Samples were acquired using BD FACS-ARIA-III™ and were analyzed using FlowJo™ software (Tree Star).

### Statistical analysis

Statistical analysis was done using GraphPad Prism Version 10.0.3 (GraphPad Software Inc). Results were analyzed using Student’s *t* -test or One-way ANOVA. Data in bar graphs are represented as mean +/- standard error of the mean (SEM). Threshold values of \**P*<0.05, \*\**P*<0.01, \*\*\**P*<0.001, or ****P<0.0001 were used to assess statistical significance.

## RESULTS

### Viral ssRNA and dsRNA mimics induce differential inflammatory and antiviral responses

Excessive inflammation caused by RNA viruses is a key determinant of poor clinical outcomes^34^. However, the key virus and host factors driving RNA virus-induced lethal inflammation and severe disease are poorly described. Since both viral ssRNA and dsRNA are released extracellularly upon lytic infection^15 35^ and are detected by the immune sensors on the sentinel or circulating myeloid cells, we evaluated whether ssRNA and dsRNA and their respective cell surface/endosomal TLRs elicit differential inflammatory cytokine and chemokine responses. For these studies, murine BMDMs were treated with ssRNA mimic (R848) and dsRNA mimic (Poly I:C). Cells collected in Trizol at 8 hours post-stimulation or cell supernatants collected at 24 hours post-stimulation were used to assess mRNA and protein levels, respectively, of inflammatory cytokines and chemokines. Our results show a significant increase in mRNA levels of inflammatory cytokines and chemokines in R848 treated BMDMs compared to Poly I:C treated cells (Figure 1A). Further examination of protein levels of representative inflammatory cytokines (TNF-α and IL-6) also revealed markedly higher levels of these inflammatory mediators in R848 treated cells compared to Poly I:C stimulated cultures (Figure 1B). Next, we examined the effect of R848 and Poly I:C on antiviral IFN-I and ISG responses. Cells in trizol and cell supernatants collected at 8 and 24 hr post-stimulation were used to measure mRNA or protein levels of IFNs and ISGs. As shown in Figure 1C-D, we observed marginal to no IFN and ISG response in R848-stimulated macrophages compared to unstimulated cells, whereas Poly I:C treatment induced a robust increase in IFN and ISG levels. Poly I:C activates both TLR3-TRIF and MDA5-MAVS in fibroblasts ^36^. Therefore, to confirm Poly I:C-induced antiviral and inflammatory response in macrophages is mediated by TLR3-TRIF signaling, we stimulated wild-type and TRIF^-/-^ cells with Poly I:C. Our results showed a complete loss of antiviral and inflammatory cytokine production in Poly I:C treated TRIF-/- cells compared to wild-type cells (Figure 1E), suggesting that Poly I:C-induced immune response in macrophages is dependent on TLR3-TRIF activity. We also confirmed that R848 mediated cytokine/chemokine response is Myd88 dependent (Figure 1F). These results collectively demonstrate that extracellularly derived viral ssRNA mimic induces significantly elevated levels of inflammatory cytokine/chemokines compared to dsRNA mimic, whereas dsRNA is a more potent inducer of IFN and ISG responses than ssRNA in mouse macrophages.

**Fig 1:**
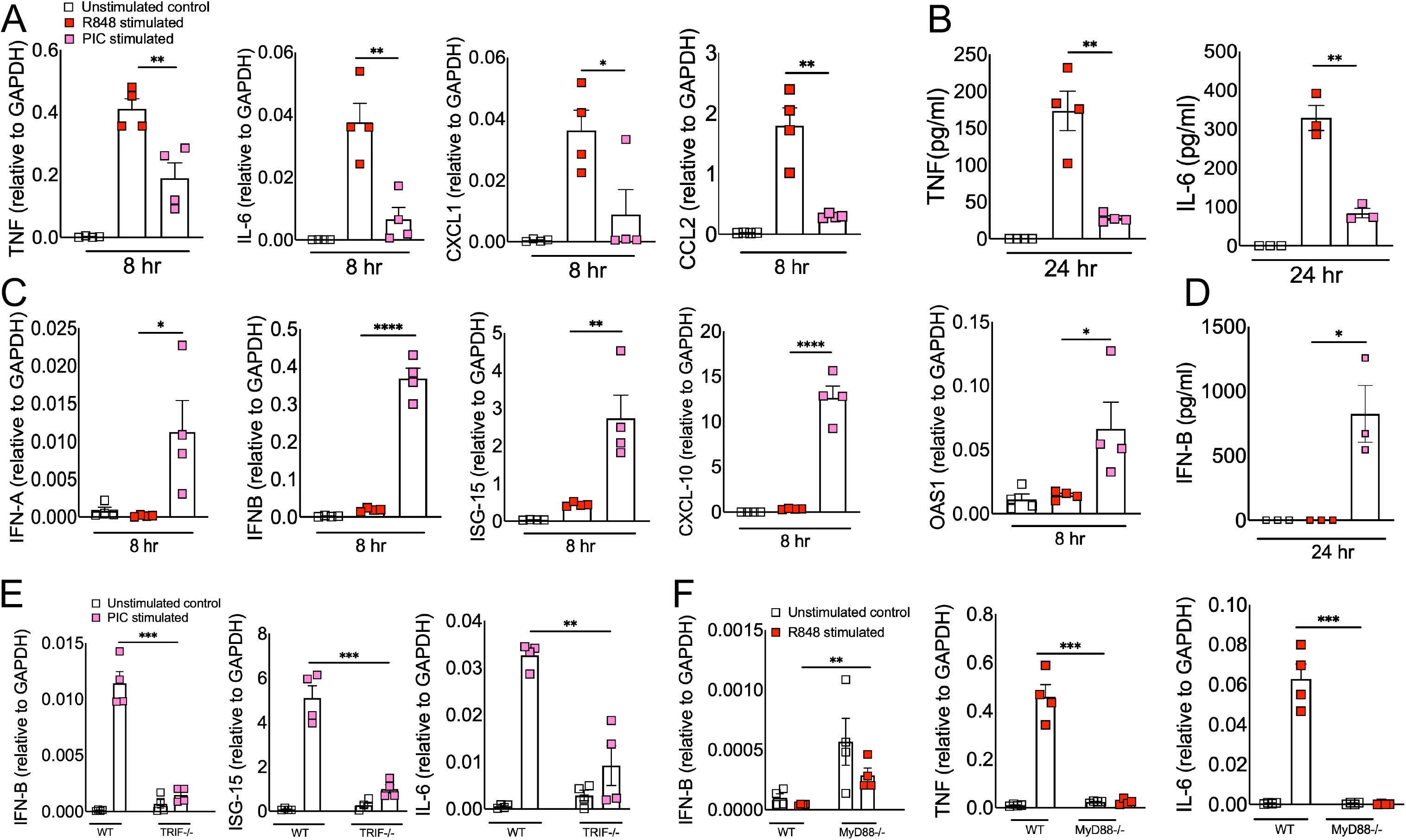
ssRNA/TLR7 stimulation induces robust inflammatory cytokine/chemokine responses whereas dsRNA mimics are potent inducers of antiviral IFN and ISG responses. Murine BMDMs were stimulated with viral RNA mimics; R848 (ssRNA mimic) and poly I:C (dsRNA mimic) each with 10 ug/ml. mRNA **(A)** and protein levels **(B)** of inflammatory cytokines and chemokines and mRNA **(C)** and protein levels **(D)** of interferons and ISGs were assessed 8 & 24 hours after stimulation respectively. WT and TRIF-/- BMDMs **(E)** and WT and MyD88-/- BMDMs **(F)** were stimulated with viral RNA mimics; poly I:C (dsRNA mimic) and R848 (ssRNA mimic) each with 10 ug/ml respectively. mRNA levels of interferons and inflammatory cytokines were assessed 8 hours post-stimulation. Data are representative of 3 independent experiments. Statistical significance was determined using Student’s t-test with *P < .05, **P < .01, **P < .001 and ****P < .0001.

### Macrophages express differential levels of cell surface and endosomal TLR3 and TLR7

Macrophages express high levels of TLRs, and recognition of viral PAMPs by these TLRs plays a crucial role in antiviral and inflammatory response ^37 38^. Extracellularly derived viral dsRNA and ssRNA are primarily recognized by cell surface/endosomal TLR3 and TLR7, respectively ^21 23^. Since we observed differential antiviral and inflammatory cytokine response following TLR3 and TLR7 agonist administration, we first asked whether altered TLR expression by macrophages contributes to differential antiviral and inflammatory cytokine production. To test this possibility, we examined cell surface and endosomal TLR3 and TLR7 expression in murine BMDMs. As shown in Figure 2A-B, flow cytometry data showed that the percentage of macrophages expressing cell surface and endosomal TLR7 was significantly higher than for TLR3 expression. Additionally, we found a 5-10-fold increase in mean fluorescence intensity (MFI, i.e., number of molecules expressed per cell) of TLR7 expression in endosomes than on the cell surface (Figure 2C). These results show a significantly higher expression of TLR7 in macrophages (both cell surface as well as intracellular) than that of TLR3, which may contribute to the differential inflammatory and antiviral immune response observed following ssRNA mimic/TLR7 agonist treatment compared to the dsRNA mimic/TLR3 agonist administration.

**Fig 2:**
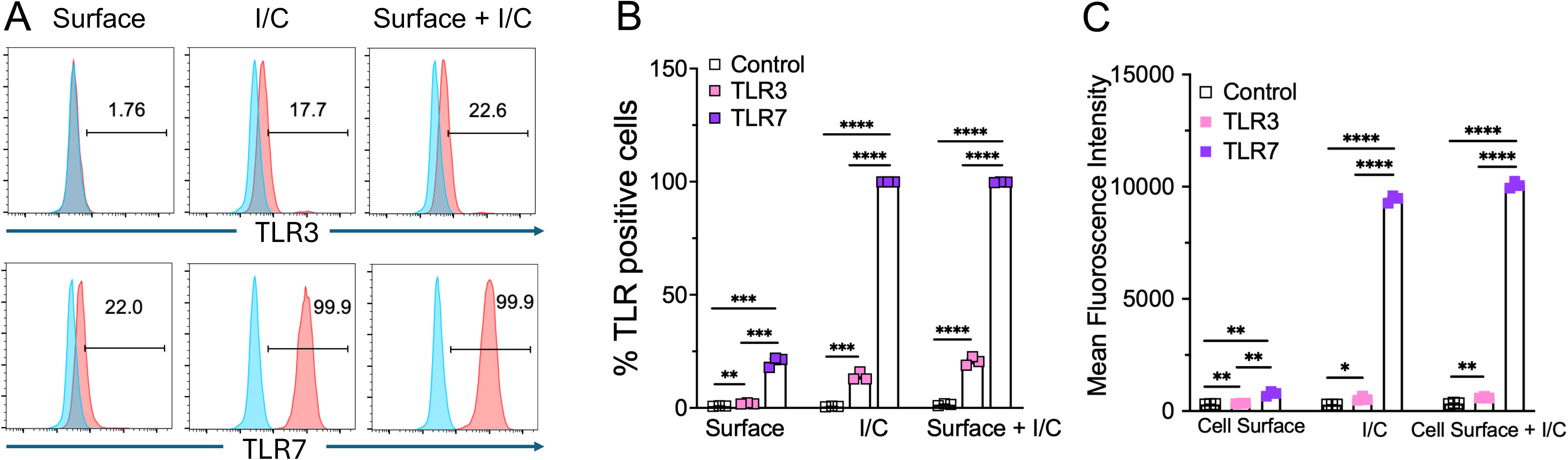
TLR3 and TLR7 expression in mouse bone marrow macrophages. Expression of cell surface and intracellular TLR3 and TLR7 in naïve mouse bone marrow macrophages was analyzed by flow cytometry (FACS) using standard protocols (**A**). Scatter plot graphs show the percentage and Mean fluorescent intensity of TLR3 and TLR7 positive cells (**B** and **C**). Data are representative of 3 independent experiments. Statistical significance was determined using Student’s t test with *P < .05,**P < .01, **P < .001 and ****P < .0001.

### R848 and Poly I:C induce differential MAPK and NF-kB activation in macrophages

Our results showed differential inflammatory cytokine response in Poly I:C and R848 treated cells (Figure 1), which correlated with cell surface and endosomal TLR3/7 expression (Figure 2). However, these results fail to show whether or not such differential TLR3/7 expression correlates with altered downstream signaling. TLR7 facilitates MyD88-TRAF6 and TLR3 utilizes TRIF-TRAF6 adapters to induce NF-kB/ MAPK-mediated inflammatory cytokine response ^39 40^. Therefore, to determine the mechanistic basis for R848/TLR7- and Poly I:C/TLR3-induced differential inflammatory cytokine response, we stimulated primary mouse BMDMs with these viral ssRNA and dsRNA mimics. The cell lysates collected at 10, 30, and 60 minutes were analyzed for total and phospho-levels of MAPKs and NF-kB using western blot assay. As shown in Figure 3A, we observed a significant increase in phosphorylation of MAPK (p-ERK, p-P38, p-JNK) and NF-kB(p-P65) pathways in R848 stimulated macrophages as early as 10mins post-stimulation, which persisted for up to 30 min post-stimulation. In contrast, we observed a delay in poly I:C induced phosphorylation of these pathways with robust MAPK/NF-kB activity observed at 60 min post-stimulation (Figure 3A-B). These results indicate that R848-TLR7 signaling promotes early activation of MAPK and NF-kB pathways, whereas poly I:C-TLR3 signaling is a delayed and muted activator of these downstream pathways.

**Fig 3:**
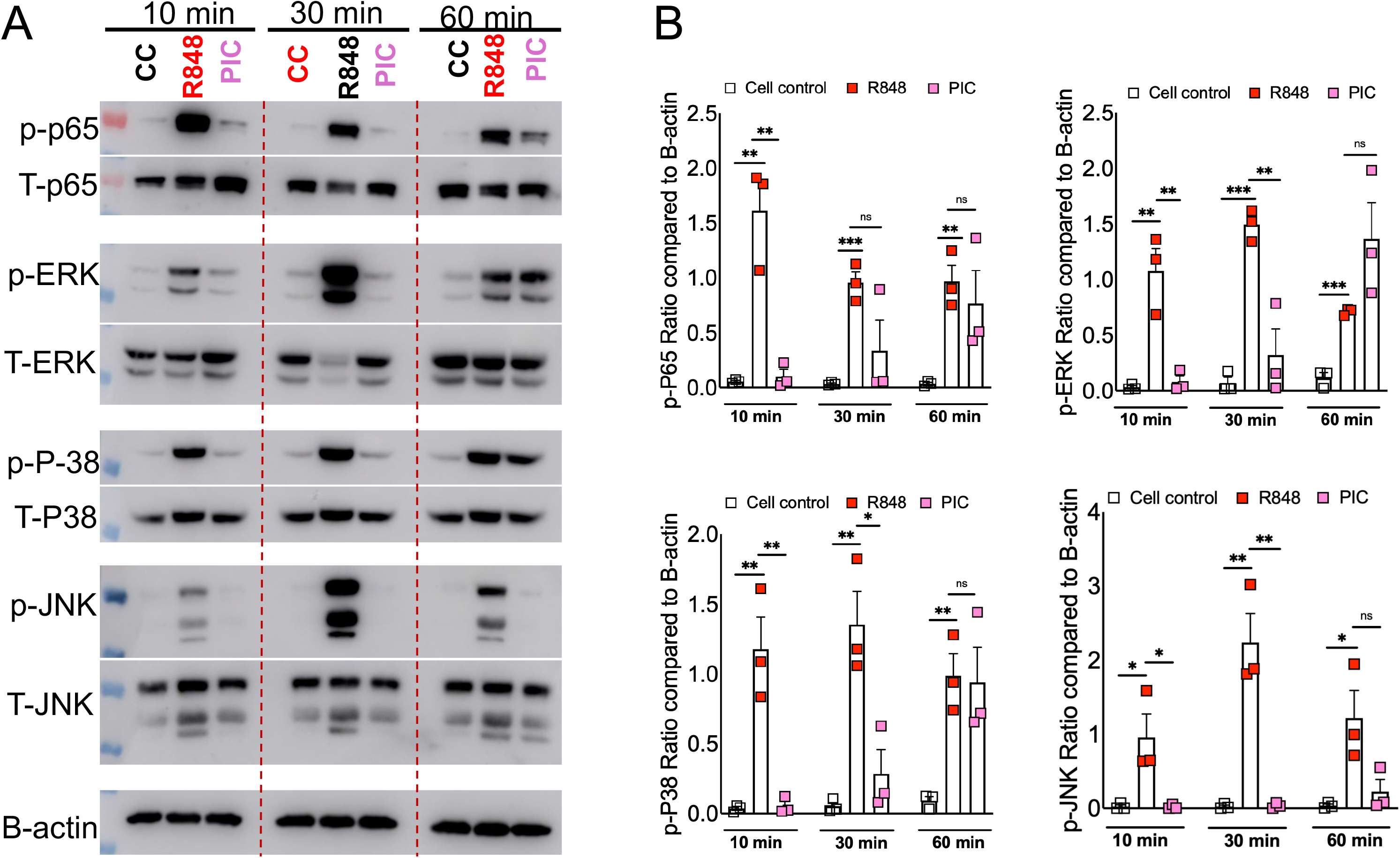
RNA mimic stimulation induced differential phosphorylation of MAPK and NF-kB signaling. Murine bone marrow macrophages were treated with ssRNA mimic (R848) and dsRNA mimic (PIC) for 10, 30, and 60 mins. Cell lysates were collected and western blot was performed using mAbs against phosphorylated MAPK (ERK1/2, JNK and p-38) and NF-kB (p-65) (**A**) and relative quantification of protein levels by densitometry analysis (**B**). Data are representative of 3 independent experiments (**A**) or pooled from 3 independent experiments (**B**). Statistical significance was determined using Student’s t-test with *P < .05, **P < .01, **P < .001 and ****P < .0001.

### Blocking ERK1/2 reduces inflammation and improves IFN in viral RNA mimic treated cells

Our results thus far illustrate that dysregulated immunity in ssRNA mimic treated cells correlates with high TLR7 expression and TLR7-induced robust MAPK/NF-kB activation (Figures 1-3). To further demonstrate that TLR7-induced robust MAPK and NF-kB activity promotes excessive inflammation, we blocked MAPK and NF-kB signaling using well-established inhibitors for each pathway. First, we evaluated the ability of MAPK and NF-kB inhibitors to block pathway-specific activity. As shown in figure 4A, TLR-stimulated BMDMs pre-treated with specific inhibitors of ERK1/2 (Trametinib), p38 (Losmapimod), JNK (JNK inhibitor I), and NF-kB (BAY11-7082) showed reduced phosphorylation of respective proteins compared to untreated cells. Next, cell supernatants collected from control and pathway-specific inhibitor-treated BMDMs stimulated with R848 or Poly I:C (10 g/ml each) were examined for TNF, MCP-1, and IFN-β levels. As expected, blocking p-ERK1/2, p-NF-kB, and p-JNK activity all reduced inflammatory cytokine and chemokine production (Figure 4B). However, to our surprise, p-p38 inhibition enhanced inflammatory cytokines/chemokine levels in BMDM cultures (Figure 4B). Interestingly, p-ERK1/2 inhibition improved type IFN-I levels (Figure 4C) in both R848 and Poly I:C treated cells, which correlated with enhanced ISG levels (Figure 4D-E). Conversely, blocking NF-kB suppressed IFN-β levels, whereas p38 and JNK inhibition had minimal impact on IFN-I response (Figure 4C). Collectively, these results establish that a) TLR7-induced robust MAPK/NF-kB activation promotes excessive inflammation and b) ERK1/2 signaling promotes inflammatory cytokine/chemokine levels while suppressing viral RNA mimics-induced IFN-I and ISG responses.

**Fig 4:**
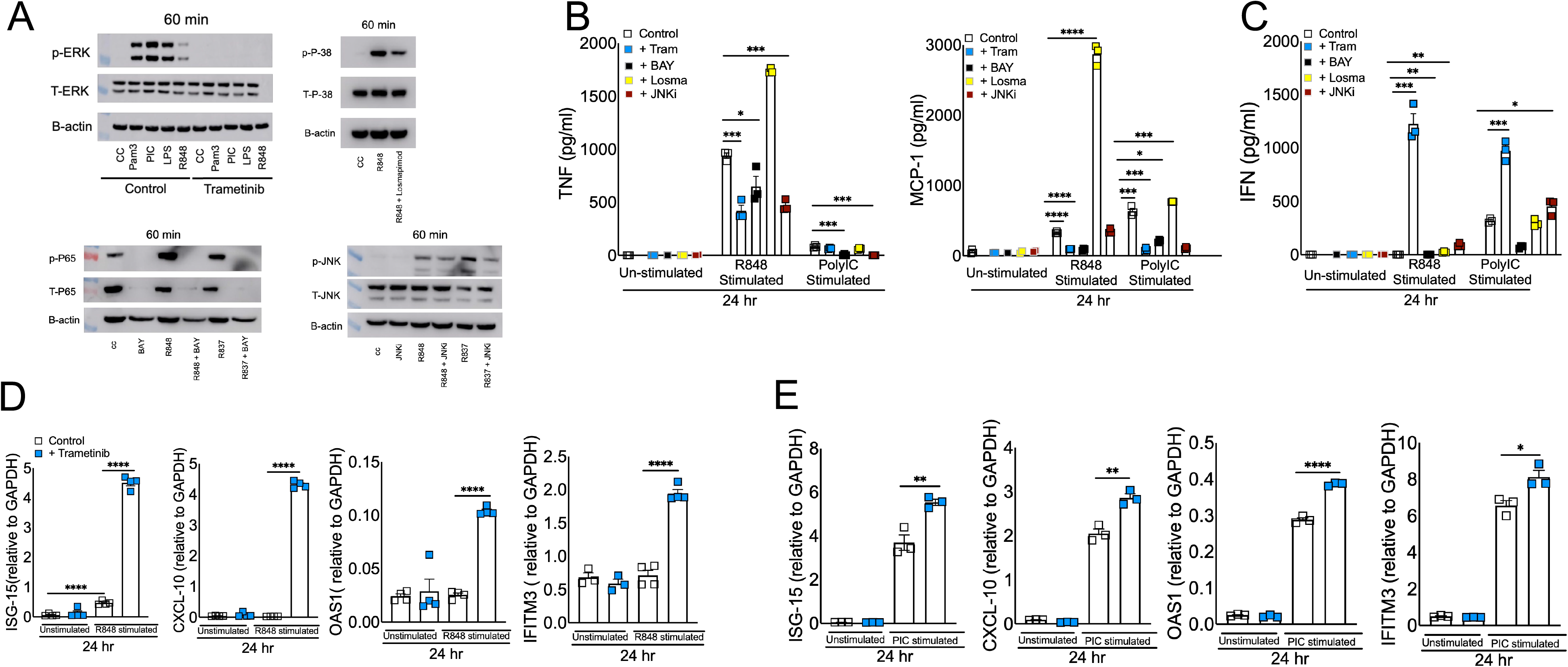
Effect of blocking MAPK and NF-kB signaling in viral RNA mimic induced inflammatory and antiviral cytokine response. Murine BMDMs were treated with each specific inhibitor of NF-kB and MAPK pathways for 2 hrs followed by stimulation with R848 and or Poly I:C each with 10 ug/ml. Western blot showing the inhibitory effect of Trametinib, Losmapimod, BAY, and JNKi on p-ERK, p-P38, p-P65, and p-JNK respectively **(A)** Protein levels of cytokines and interferons (**B and C**) and mRNA levels of ISGs **(D and E)** were assessed at 24 hours post-stimulation. Data are representative of 2-3 independent experiments (**B-E)**. Statistical significance was determined using Student’s t-test with *P < .05, **P < .01, **P < .001 and ****P < .0001.

## DISCUSSION

Emerging and re-emerging RNA viruses cause acute and often fatal illness. The RNA virus-induced severe disease is caused by excessive inflammatory and impaired or delayed antiviral IFN/ISG responses ^9 10^. Additionally, ssRNA-induced severe inflammation is associated with excessive monocyte-macrophage accumulation ^41 42 43 44^. However, the basis for RNA virus-induced lethal inflammation and the host factors that suppress antiviral IFN/ISG responses need to be well defined. Viral RNAs are critical orchestrators of antiviral and inflammatory responses and monocyte-macrophages are a key source of inflammatory mediators and IFNs ^45 8 46^. Therefore, in this study, using RNA mimics, we examined the effects of viral ssRNA and dsRNA in eliciting macrophage-induced inflammatory and antiviral responses. Our results illustrate that viral ssRNA mimics induce a robust inflammatory but a poor IFN response, whereas dsRNA mimics elicited a strong IFN and a relatively weak inflammatory cytokine response. We further demonstrate that viral ssRNA- and dsRNA-mimic-mediated differential immune response is likely driven by the high cell surface and endosomal TLR7 expression as opposed to low TLR3 expression in both compartments. Excessive inflammatory cytokine response and high cell surface/endosomal TLR expression correlate with early and robust NF-kB and MAPK activation in TLR7 compared to TLR3 agonist-treated cells. Finally, we demonstrate that blocking NF-kB and MAPK sub-pathways reduces viral RNA mimic-induced inflammatory cytokine production, while only ERK1/2 inhibition significantly increased IFN/ISG responses. Our results, albeit using viral RNA mimics, collectively explain the basis for the dysregulated immunity caused by RNA viruses.

The differential role of TLR7 and TLR3 in inducing inflammatory and antiviral cytokines have been previously described ^21 47 48^. However, most of these studies have examined either TLR7 and TLR3 agonist-induced inflammatory cytokine response or antiviral IFN/ISG response without defining the implications of these results in RNA virus-induced dysregulated immunity. The majority of emerging RNA viruses consist of ssRNA genome that make dsRNA intermediate during replication ^49^. ssRNA is 100-fold or more abundant than dsRNA during active ssRNA virus replication ^16^. Therefore, we believe that it is critical to clearly define the role of viral ssRNA and dsRNA in eliciting inflammatory cytokine/chemokine and antiviral immune responses. In this study, we used well-described viral ssRNA and dsRNA mimics to examine both mRNA and protein levels of inflammatory cytokines and IFN/ISGs at both early and late times to comprehensively describe the role of viral RNAs in monocyte-macrophage mediated immune response. Although the use of viral ssRNA and dsRNA mimics instead of authentic viral ssRNA or dsRNA is a limitation in our study, we strongly believe that our results will lay a foundation for a careful examination of virion purified, invitro synthesized, or in-vivo generated viral ssRNA and dsRNA in causing dysregulated immunity and severe disease.

In addition to detailed examination of viral ssRNA and dsRNA mimic-induced inflammatory and antiviral immunity, we provided a potential mechanistic explanation for differential immune response induced by viral RNA mimics. We illustrate that high cell surface and endosomal TLR7 expression compared to TLR3 levels correlate with R848-induced robust inflammatory cytokine production. Conversely, cell surface/ endosomal TLR7 levels did not correlate with IFN response, as shown by low IFN/ISG levels in R848-treated cells compared to Poly I:C-treated macrophages. These results suggest TLR7/Myd88 signaling is a weak IFN-I/ISG inducer compared to TLR3/TRIF activity. An additional explanation for Poly I:C-induced high IFN/ISG levels is that the dsRNA is escorted into the cytosol and sensed by MDA5 ^50 17^. However, our results show that at least in macrophages, Poly I:C-induced IFN/ISG response is mediated via TLR3-TRIF pathway (Figure 1E). We further show that high TLR7 and inflammatory cytokine levels correlate with robust NF-kB and MAPK (p38, ERK1/2, and JNK) activity as early as 10 minutes post-R848/TLR7 stimulation compared to Poly I:C treatment (Figure 3). Additionally, TLR7-induced robust activation of NF-kB and MAPK at both 30- and 60-min post-stimulation (Figure 3). These results collectively suggest that timing and strength of NF-kB and MAPK activity dictate the level of inflammatory cytokine and chemokine production in viral RNA mimic treated cells. Although previous studies have examined the role of either NF-kB or MAPK activity in viral RNA mimic treated and RNA virus-infected cells, our work comprehensively examines multiple key signaling pathways at several time points providing critical and detailed insight into the role played by these pathways in viral RNA induced inflammation and antiviral response ^51 52^.

We show robust activation of multiple pathways in ssRNA mimic treated cells and identify the impact of blocking these pathways in mitigating excessive inflammation. Significant reduction in inflammatory cytokine/chemokine levels in NF-kB-, ERK1/2-, and JNK-inhibitor-treated cells highlights the importance of simultaneously blocking multiple pathways to mitigate viral ssRNA-induced inflammation. Of note, in contrast to published results demonstrating the anti-inflammatory functions of FDA-approved p38-MAPK inhibitor Losmapimod, we consistently observed a significant increase in inflammatory cytokine/ chemokine levels in Losmapimod treated TLR7 stimulated, but not Poly I:C treated macrophages ^53 54^. These results suggest that blocking p38 likely exacerbates viral ssRNA-induced inflammation. Although it is unclear if such response is specific to Losmapimod treatment in BMDMs or is a general response to p38 inhibition in all cell types, these outcomes warrant a further investigation of immunomodulatory effect of Losmapimod, particularly during RNA virus infection. Another important finding from our study is that unlike blocking NF-kB and JNK signaling, ERK1/2 inhibition significantly improved antiviral IFN/ISG response in ssRNA and dsRNA mimic treated cells, emphasizing the importance of blocking ERK1/2 activity to suppress excessive inflammatory cytokines and simultaneously improve antiviral IFN/ISG response. Although corticosteroids are currently used to suppress exuberant inflammation, corticosteroid-mediated suppression of antiviral innate and adaptive immunity is a clinical concern ^55 56^. Therefore, blocking ERK1/2 signaling along with NF-kB and other MAPK pathways is likely superior to provide better protection compared to the current immunomodulatory therapies.

In summary, using viral RNA mimics, we demonstrate that ssRNA, which is abundantly produced during RNA virus infections, is potentially a key mediator of virus-induced exuberant inflammation and impaired antiviral IFN/ISG response. Mechanistically, we show that high cell surface and endosomal TLR7 expression drives robust NF-kB and MAPK activation, thus facilitating excessive inflammatory cytokine response. Additionally, given ERK1/2 signaling suppresses antiviral response and promotes inflammation, blocking ERK1/2 activity is perhaps a better immunomodulatory strategy to mitigate dysregulated immunity and provide better protection.

## Author Contributions

RC and RS conceptualized the study; RS, PJ, RG, and CW carried out experiments and data analysis; RC and RS wrote the manuscript; RC, RS, PJ, RG, and CW reviewed and edited the manuscript. RC provided mentoring and supervised experiments, data analysis, and presentation.

## Acknowledgments

This work is supported in part by an institutional research fund to RC from Oklahoma State University, College of Veterinary Medicine, NIH R21AI186028, Oklahoma Center for Respiratory and Infectious Disease (OCRID) Centers of Biomedical Research Excellence (COBRE) grant NIH P20GM103648 and NIHAG060222. We also acknowledge OCRID/CoBRE Immunopathology Core for assistance flow cytometry studies.

